# Boosting forward connectivity between primary visual and body selective cortex reduces interference between sex and emotion judgements of bodies

**DOI:** 10.1101/2024.09.10.612296

**Authors:** Marco Gandolfo, Giulia D’Argenio, Paul E. Downing, Cosimo Urgesi

## Abstract

We effortlessly categorise other people along socially relevant categories such as sex, age, and emotion. A core question in social vision relates to whether we perceive these categories independently or in relation to each other. Here, we investigated categorisation of sex and emotion from the body, finding that participants generally fail to fully ignore task-irrelevant variations of sex while judging body emotional expressions. In contrast, sex categorisation was unaffected by variations in emotional expression. This asymmetric interaction between sex and emotion may arise because of bottom-up visual processing, due to partially shared visual features used for both judgments, or because of top-down, categorical associations between sex and emotion categories. To disentangle these possibilities, we used cortico-cortical paired associative stimulation (ccPAS) to modulate the connectivity between primary visual cortex and the extrastriate body area. We posited that boosting forward connectivity between these regions would increase efficiency of feature-based processing, while boosting feedback connectivity would enhance the separability of semantic categories related to sex and emotion. We found that boosting forward connectivity eliminated the interference of sex on emotion judgments, while that interference remained unaffected with modulation of feedback connectivity. These findings suggest that interactions between sex and emotion in body perception emerge during the perceptual analysis of the stimuli, and add to our understanding of person perception and social categorization.

## 1. Introduction

Observers tend to make spontaneous categorical attributions about other people based upon their appearance (R. B. Jr. Adams et al., 2011; Liberman et al., 2017). This rapid process engages semantic knowledge and stereotypes associated with social groups such as race, sex^1^, or age (Fiske & Neuberg, 1990) that also intersect with the evaluation of current states (such as direction of attention or emotional expression; Bijlstra et al., 2019; Hedgecoth et al., 2023). A core question concerns the *automaticity* of categorisation from visual appearance: to what extent does the processing of category cues depend on other task demands and the observer’s goals? Recent dynamic views of social perception posit a continuous interaction between visual bottom-up and semantic top-down social-cognitive processes in the interpretation of social stimuli (Freeman & Ambady, 2011; Freeman & Johnson, 2016; Stolier & Freeman, 2016). In this conception, visual features influence the competition between potential categorical interpretations, while simultaneously task goals, stereotypes, and semantic associations bias the selection and perception of visual features. Building on this framework, here we used a novel neurostimulation approach to examine top-down and bottom-up processes in the case of categorizing *sex* and *emotion* from images of the human body. Before expanding on the logic of this pairing of social dimensions, we first review some of the relevant background on automaticity of social vision.

Researchers have identified several senses of the umbrella term “automaticity’ (Bargh, 1989). One of these, our focus here, describes a cognitive process that unfolds similarly whether or not it aligns with the observer’s goals. For example, do we automatically infer the ethnicity of another person, even when our focus is specifically on their emotional expression? A powerful approach to testing this sense of automaticity was developed by Garner and colleagues (Garner, 1974; Algom & Fitousi, 2016; Burns, 2016). Participants make binary judgments about a series of stimuli (e.g. on shape: square vs circle). In an *orthogonal* condition, those stimuli vary orthogonally along a task-irrelevant dimension at the same time (e.g. color: red vs green). Performance is compared to *control* conditions in which the task-irrelevant dimension (e.g. colour in the shape task) is held fixed at one level (e.g. green) while the task-relevant dimension varies. If variation on the irrelevant dimension can be successfully ignored (i.e. equivalent performance in the two conditions), this implies successful filtering by attention. That is, the irrelevant dimension is not being automatically processed during attention to the relevant one. Alternatively, if performance is hindered by irrelevant variation, this implies some degree of automatic processing of that dimension.

Early studies (Garner, 1974, 1978) applied this approach to basic aspects of perceptual processing, such as shape, texture, size. More recently, the same logic has been applied to more complex and meaningful stimuli (Cant & Goodale, 2009; Gandolfo & Downing, 2020b; Johnstone & Downing, 2017; Tharmaratnam et al., 2021). Several studies used the Garner paradigm to test predictions from neurocognitive models of face perception (Atkinson et al., 2005; Becker, 2017; Ganel & Goshen-Gottstein, 2002, 2004; Karnadewi & Lipp, 2011; Le Gal & Bruce, 2002; Schweinberger et al., 1999; Schweinberger & Soukup, 1998). Such models often posit a distinction between dynamic properties (e.g. momentary emotional expressions) and static properties (e.g. shape and texture features signaling identity or sex) of the face (Bruce & Young, 1986; Duchaine & Yovel, 2015; Haxby et al., 2000). However, evidence from Garner tasks has generally (although not always) pointed to interactions between processing of static and dynamic aspects of the face, rather than full independence between these two types of information.

For example, researchers investigated the relationship between sex (a relatively stable characteristic of the face) and emotional expressions (a changeable aspect of the face). Using the Garner logic, they showed asymmetrical Garner interference (Atkinson et al., 2005; Karnadewi & Lipp, 2011; but see Le Gal & Bruce, 2002). Participants could not fully ignore irrelevant variation of sex while categorizing the emotional expression, but they could generally ignore emotional expression while judging sex. This suggests that more stable visual cues (i.e. those related to sex dimorphisms in the face) are harder to ignore because they provide a reference for computation of more changeable social cues such as emotional expressions. These findings illustrate the usefulness of the Garner logic for understanding categorisation of complex meaningful stimuli such as the face.

The shape and posture of the body, as well as the face, convey meaningful signals about the states and traits of other individuals (e.g. emotions: Aviezer et al., 2012; de Gelder, 2009; identity: Rice et al., 2013; Yovel & O’Toole, 2016; sex and gender: D’Argenio et al., 2020, 2022; Gandolfo & Downing, 2020a; Johnson & Tassinary, 2005; postures: Han et al., 2024, 2025; Reed et al., 2006). Some studies have applied the Garner logic to investigate whether these signals are automatically or independently processed (Craig & Lipp, 2023; Gandolfo & Downing, 2020b; Johnstone & Downing, 2017; see also Reed et al., 2018; Bijlstra et al., 2019). Of particular interest for the present study is previous work examining the perception of sex and emotional expressions from the body. The relationship between sex and emotion provides a useful test case, because it provides another test of the distinction between dynamic versus stable characteristics described above, complementing the larger evidence base from faces. Finally, the existence of well-characterised body-selective brain regions of interest provides suitable targets for a neurostimulation approach (Peelen & Downing, 2007, 2017).

To date, there have been two directly relevant studies (Gandolfo & Downing, 2020b; Craigg & Lipp, 2023). Gandolfo and Downing (2020) reported evidence for independence between sex and emotional expression. Across three experiments using angry and sad bodies, angry and happy bodies, and happy and sad bodies, they found no evidence of Garner interference (supported by Bayesian analyses indicating strong support for the null effect). Participants could report either emotional expression or sex, without performance being affected from irrelevant variation of the other cue. Conversely, Craig and Lipp (2023), using a different stimulus set (BESST, Thoma et al., 2013), reported symmetrical Garner interference in one experiment using angry and happy body expressions, and no Garner interference between sex and emotion when using angry and sad body expressions. These results pointed to a reciprocal influence of sex and emotion processing, at least when the emotion categorization involved emotions with different valence. The effect size, however, was relatively greater for the emotion task, suggesting greater influence of sex processing on emotion processing than *vice versa*.

This mixed pattern of results raises questions about when cues to sex and emotion may interact and at what level. Such interactions may emerge through a bottom-up perceptual process (see also Jugović et al., 2024), for example when the typical visual features of an angry expression are more similar to typically masculine than feminine shape (Becker et al., 2007). Additionally, top-down processes may contribute. For example, stereotypes at a semantic level may associate femininity with positive rather than with negative emotional states (Bijlstra et al., 2019; see Craig & Lee, 2020 for a review). In the former case, Garner interference between sex and emotion would be due to objective covariation of the visual features that provide cues for judgments of emotional expressions and sex. In the latter case, an interaction would be the due to feedback processing of social categories influencing how we interpret visual information about the body.

Neuroscientific evidence may help to supplement previous behavioural evidence on distinguishing top-down and bottom-up processes in the categorisation of social stimuli. Specifically, here we adopt the extrastriate body area (EBA) and primary visual cortex (V1) as a simplified model system to examine feedforward and feedback connections between early perceptual processing of stimulus features, and higher level processing of category-specific object properties. EBA is a focal occipitotemporal region that has been shown with fMRI to respond strongly and selectively to images of human bodies depicted in a range of image formats (Downing et al., 2001; Peelen & Downing, 2005; Gandolfo et al., 2024). Transcranial magnetic stimulation (TMS) investigations confirm its category-specific functional role in detecting and discriminating bodies and body parts (Candidi et al., 2008; Gandolfo et al., 2024; Gandolfo & Downing, 2019; Pitcher et al., 2009; Urgesi et al., 2004, 2007; van Koningsbruggen et al., 2013; see Peelen & Downing, 2017 for a review).

We adopt the ccPAS protocol of TMS (cortico-cortical paired associative stimulation; (Koch et al., 2013; Rizzo et al., 2009; for reviews see Tarasi et al., 2024; Di Luzio et al., 2024; Hernandez-Pavon et al., 2023) to assess whether strengthening feed-forward or feed-back connectivity between V1 and EBA modulates the ability to filter out task-irrelevant information during categorization of sex and emotion from the body. ccPAS is a dual-site offline stimulation protocol in which pairs of TMS pulses are repeatedly applied to two interconnected cortical areas at a specific and constant inter-stimulation interval (ISI). ccPAS is used to modulate connectivity and it is based on Hebbian principles (Hebb, 1949; Caporale & Dan, 2008). According to these principles, synapses are potentiated when presynaptic neurons coherently and repeatedly fire before postsynaptic neurons. The ccPAS protocol is considered to increase the synaptic efficacy of the connections between two target areas, showing long-term potentiation-like effects (Buch et al., 2011; Koch et al., 2013; Romei et al., 2016; Santarnecchi et al., 2018). The ccPAS protocol has been previously applied to modulate connectivity in visual cortex (Borgomaneri et al., 2023; Chiappini et al., 2018, 2024; Di Luzio et al., 2022; Romei et al., 2016). In a seminal study, for example, ccPAS stimulation of reentrant projections from visual motion-selective V5 to V1 was shown to enhance visual sensitivity to motion by lowering the threshold for motion coherence discrimination (Romei et al., 2016).

As a conceptual replication of previous work, we first tested with the Garner paradigm whether participants could selectively attend to either sex (male/female) or emotion (happy/fearful) in series of body images, while filtering out the other, irrelevant dimension. Participants’ behavioural performance revealed asymmetrical interference: they could not fully ignore irrelevant variation of sex while categorizing emotional body expressions, but they were able to ignore emotional expression while judging sex.

Next, in a second experiment, using cc-PAS we found that the interference of body sex on emotional expression judgments was eliminated when boosting forward connections from V1 to EBA, but not when stimulation targeted connectivity in the opposite direction, from EBA to V1. This pattern implies that interactions in processing sex and emotions from the body arise at a forward stage of processing due to the bottom-up salience of sex cues. In brief, our claim is that by enhancing the efficiency of feed-forward connections from primary visual to body-selective visual cortex, we enhanced processing of those specific visual cues that are relevant for the task at hand, thus eliminating the interference of sex on emotion judgements.

## 2. General Methods

### 2.1 Participants

Thirty-two participants took part in the behavior-only experiment (Experiment 1; 30 women, 21 +/- 3 years). Four participants declared themselves to be left-handed. All the participants were Education students at the University of Udine who participated in return for course credit. This sample size allows detecting with 80% power an effect of medium size (dz = 0.51). In addition, we chose this sample size to match previous research using this paradigm (Gandolfo and Downing, 2020; Johnstone and Downing, 2017).

Twenty-nine different participants took part in the TMS experiment (Experiment 2; 23 women, 23 +/- 2 years). Participants were students at University of Udine who participated in return for course credit. They were screened following the safety screening standard questionnaire for rTMS (Rossi et al., 2009, 2011). None of the participants reported any history of neurological, psychiatric, or other major medical disorders. All the participants reported themselves to be right-handed. We aimed to match the sample size of the behavior-only experiment. However, due to participants dropping out after registration for the study, we were able to reach a sample size of 29 participants.

For both experiments, the procedures were approved by the local Ethics Committee (Institutional Review Board, Department of Language and Literature, Communication, Education and Society, University of Udine, Italy; CGPER-2019-12-09-03) and were carried out in accordance with the ethical standards of the Declaration of Helsinki. All participants were naïve to the purposes of the experiment and gave written informed consent for their participation.

### 2.2 Stimuli

Examples of the stimuli are shown in **Figure 1A**. In Experiment 1, the stimuli were selected from two different stimulus sets used in previous studies that investigated the perception of sex and emotional expressions from the body. Stimulus set 1 was drawn from Gandolfo and Downing (2020). Stimulus set 2 was drawn from Borgomaneri et al. (2012). From each of these stimulus sets we selected 4 male and 4 female models, each expressing a fearful and a happy expression, for a total of 16 photographs per stimulus set. We chose these two emotions because these hadn’t been tested in combination in Gandolfo and Downing (2020), and because they were most similar with each other in the emotional intensity they depicted. For Experiment 2, we used only the Borgomaneri et al. (2012) stimulus set because 1) it showed a more reliable pattern of Garner interference in Experiment 1, and 2) it did not show a difference at baseline between the two tasks (see the Results section). All the images were converted to grayscale and resized to 400 (w) x 600 (h) pixels.

**Figure 1.**
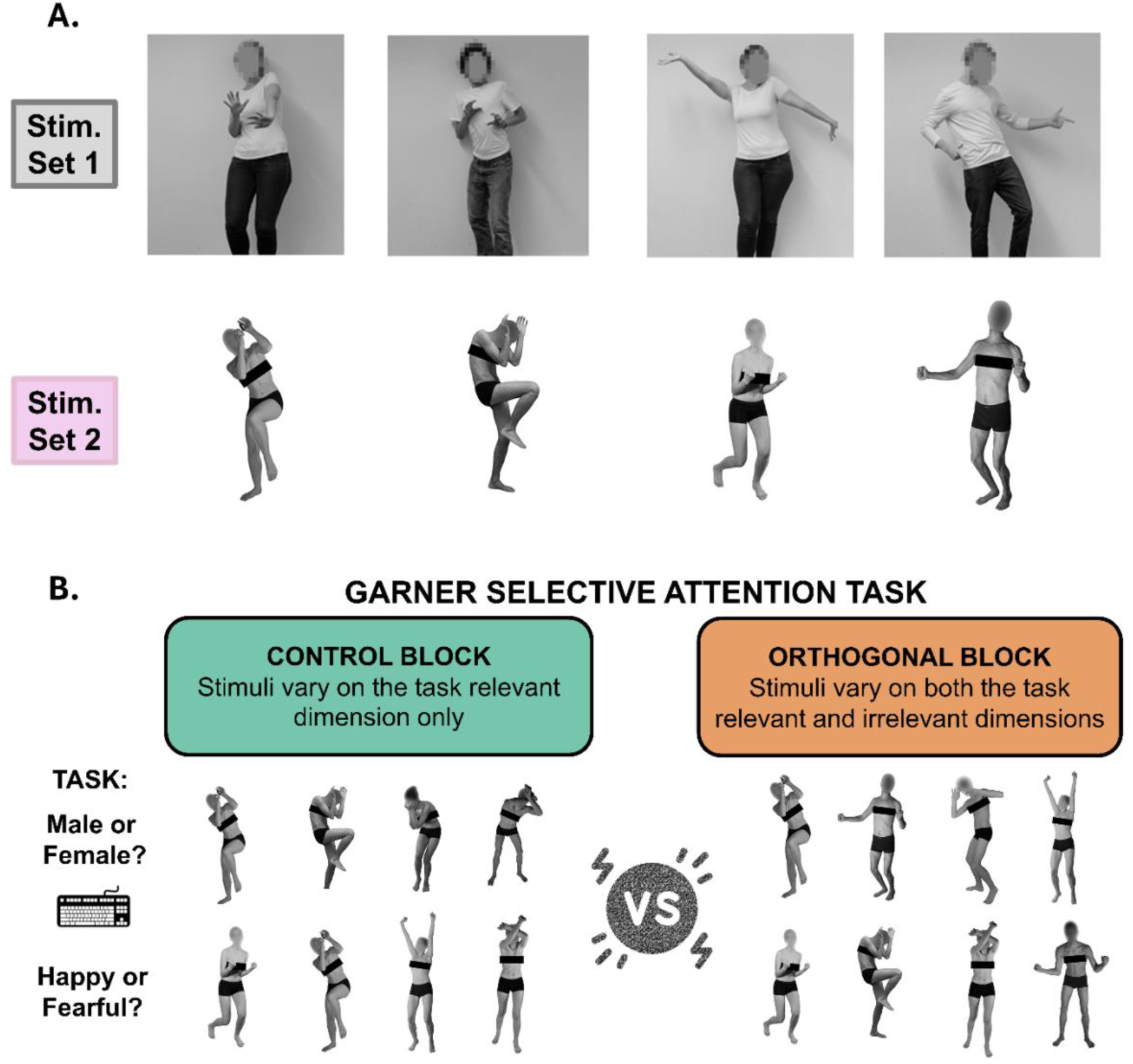
**A.** *Examples of the two stimuli sets used for Experiment 1. Row-wise they show a fearful female, fearful male, happy female, and happy male example for stimulus set 1 (Gandolfo and Downing, 2020) and stimulus set 2 (Borgomaneri et al., 2012).* **B.** *Schematic of the Garner selective attention task. Performance is compared on accurate response times between the control and orthogonal blocks. In the control blocks, the stimuli vary only on the task relevant dimension – e.g. male and female in the first left row, all expressing fearful emotions; happy and fearful in the bottom left row, all females. In the orthogonal blocks the stimuli vary on both their dimensions. Garner interference refers to increased response times in the orthogonal with respect to the control blocks. The presence of Garner interference suggests that participants cannot fully ignore the variation of the task irrelevant dimension, leading to a cost on the behavioural performance.*

### 2.3 Design

In Experiment 1, 16 participants first performed an emotion categorisation task (Happy vs Fearful), followed by a sex categorisation task; the other 16 performed the tasks in the opposite order. Task order was assigned on the basis of registration for the study. Each participant performed 1024 trials, 512 for the Borgomaneri et al. (2012) stimulus set, and 512 trials using the Gandolfo and Downing (2020) stimulus set. The two stimuli sets were presented in different macro-blocks interrupted by a 5-minute break and counterbalanced across participants.

To counteract carryover effects, for each task, Control and Orthogonal blocks were presented in a counterbalanced order across participants (see **Figure 1B**). Both the Control and the Orthogonal block comprised 128 trials. However, while both the task irrelevant and the task relevant dimensions randomly varied within the Orthogonal block, the Control block was divided in two 64-trial halves, to include trials for each level of the irrelevant dimension, which remained constant within each half. The order of the two levels of the irrelevant dimensions was also counterbalanced between participants. For example, in the Happy vs Fearful task, in one half of the Control block the images were all of females, and in the other all males. Continuing the example, in the Orthogonal block, the images would be a mixture of males and females, each displaying each emotion equally often. Block structure was not made explicit to the participants, in order to avoid drawing their attention to changes in the irrelevant dimension. The total number of trials was 256 per task, per stimulus set. Participants had a break between the two categorization tasks.

In Experiment 2, all the details were as for Experiment 1, except that we only used the Borgomaneri et al. (2012) stimulus set, and participants completed the task twice, once after each of two stimulation sessions, for a total of 1024 trials. In each session, 15 participants started with the emotion task and 14 participants with the sex task.

### 2.4 Procedure

Two speeded binary classification tasks were performed by each participant (see **Figure 2A**). In the emotion task, participants had to categorise the depicted emotional expression (fearful or happy) via key presses (f or j). In the sex task, participants had to categorise the sex (male or female), again via key presses (f or j). The key-response association was counterbalanced across participants. Each trial started with a fixation cross with a random duration between 500 to 1200 ms. Next, the body image appeared on screen for 250 ms. Following the image, the screen remained blank for 1250 ms or until the participant responded. Participants were instructed to respond as quickly as possible without sacrificing accuracy. Importantly, in the beginning of the experiment participants were not told anything about the irrelevant dimension. For example, if a participant started with the emotion task, no mention was made about the relevance of sex until they began the sex categorisation task.

**Figure 2.**
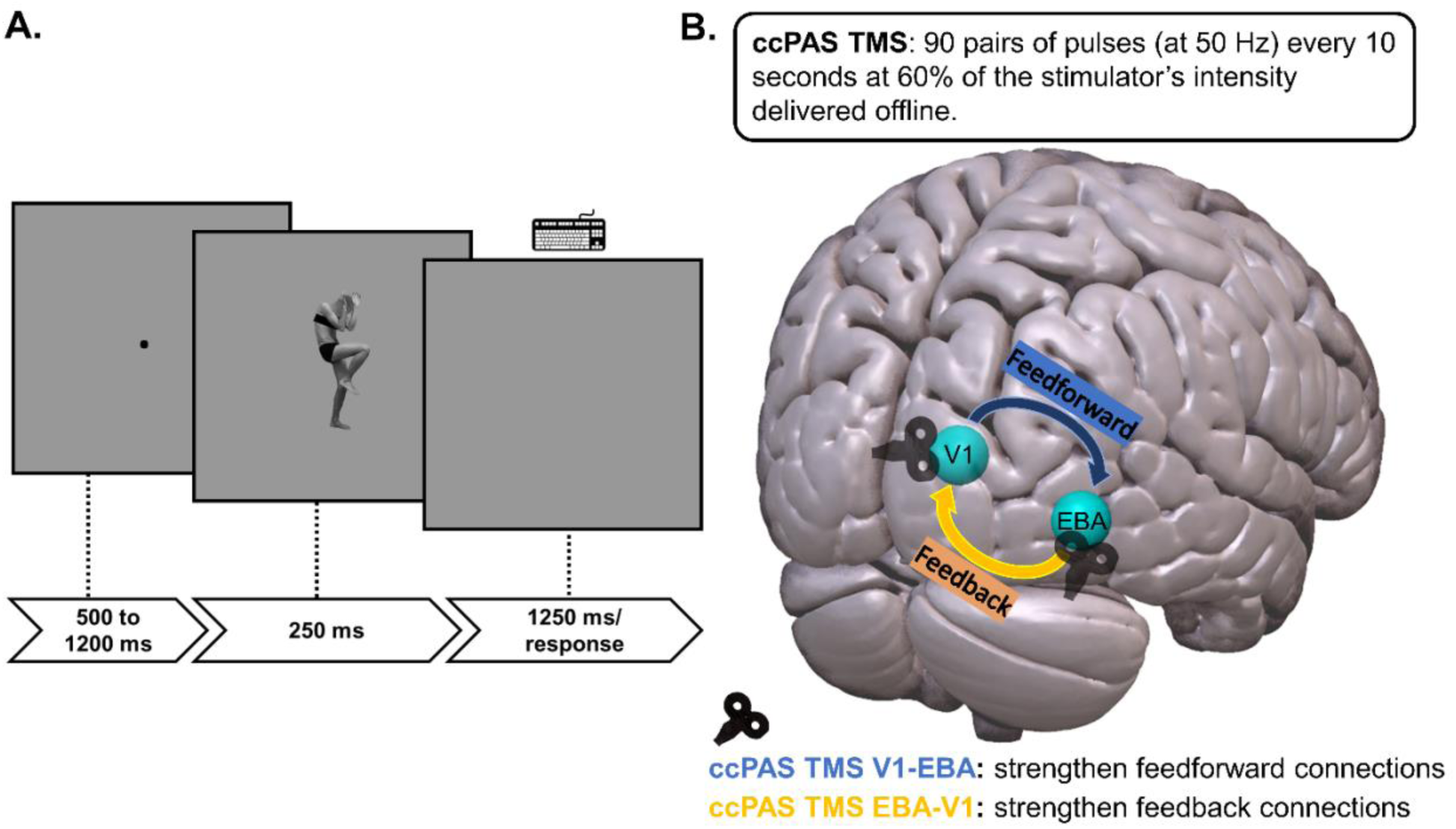
**A.** *Timeline of the task procedure used for Experiment 1 and Experiment 2.* **B.** *Graphical representation of the brain regions stimulated with TMS in Experiment 2. ccPAS TMS was delivered offline to influence feedforward or feedback connections between body-selective EBA and the primary visual cortex (V1).*

The task procedures were the same for both experiments. Experiment 1 was administered in a silent university office room, Experiment 2 in a lab environment in the presence of the two experimenters (MG and GDA). Both experiments used a 21-inch LCD screen with full HD (1920×1080px) resolution. The experiments were administered using OpenSesame (Version 3.2.7, Mathôt et al., 2012) for Windows (Microsoft, Inc.).

### 2.5 ccPAS TMS protocol and stimulation procedures

There were two stimulation sessions for each participant in Experiment 2: V1-EBA, to strengthen feed-forward connectivity between visual cortex and body selective cortex, and EBA-V1, to strengthen feedback connectivity from body selective cortex to early visual cortex. All TMS stimulation was performed over the right hemisphere. We stimulated the right hemisphere because visual body-related effects have been consistently shown on the right hemisphere (Gandolfo & Downing, 2019; van Koningsbruggen et al., 2013; Urgesi et al., 2004, 2007; Vangeneugden et al., 2014; but see Gandolfo et al., 2024).

We used a Magstim BiStim^2^ TMS stimulator (Magstim; Whitland, UK) with two Magstim figure-8, binding-iron 50mm coils. Stimulation intensity was set at 60% of the maximum stimulator output. TMS was delivered offline, 30 minutes before the participants performed the task. In each stimulation session, 90 pairs of stimuli with an ISI of 20ms were continuously delivered at a rate of 0.1 Hz (a pair of pulses every 10 seconds) for ∼15 minutes. These stimulation parameters were chosen based on a previous ccPAS TMS study targeting the primary visual cortex and V5 to investigate visual motion perception (Romei et al., 2016). We used that study as a benchmark for the present work due to the anatomical proximity between EBA and V5 (Downing et al., 2001; Weiner & Grill-Spector, 2011).

The TMS pulses were triggered remotely by sending a TTL pulse to the stimulator using MATLAB (Version 8). To target V1, the coil was centered 2cm dorsal to the inion, corresponding to the scalp position where phosphenes in the center of the visual field are usually elicited (Salminen-Vaparanta et al., 2014). For the right hemisphere EBA, we targeted the Tailarach coordinate x = 52 y = -72 z = 4 (Calvo-Merino et al., 2010; Urgesi et al., 2007) on an individually-adapted MRI template-based model obtained with an infrared neuronavigation system (Softaxic Optic, Version 3.0). Using the same reconstruction, we also registered the coordinate of V1 and double-checked whether this fell on the primary visual cortex (Brodmann Area 17) of the participant’s estimated brain image. Whilst performing the stimulation, the coordinate of of the EBA stimulation site was monitored online.

Stimulation was performed by two experimenters (MG and GDA), one holding each coil. The coil weight was further supported by compatible tripods. For the EBA, the handle was held at an angle of 120 degrees clockwise tangentially to the scalp (Urgesi et al., 2004; Gandolfo & Downing, 2019; Gandolfo et al., 2024); for V1, it was held at 180 degrees.

During stimulation, the participants rested their head on a chin-rest and fixated a cross on a gray background on a monitor screen. Based on Romei et al. (2016), effects of ccPAS stimulation in visual cortex were expected between 30 and 90 minutes after the stimulation. Therefore, we asked participants to wait 30 minutes after each stimulation session before doing the task. During the time between the stimulation and the task, we asked participants to rest while sitting on a chair or laying down, and to avoid eating food or using their mobile phone unless strictly necessary. The two stimulation sessions were separated by a 90-minute break to reduce chances of carry-over effects of one stimulation session to the other.

### 2.6 Analyses

In both experiments we considered as outliers participants who performed (in either accuracy or reaction times) 3 SDs above or below the group mean (averaging across all conditions). Based on these criteria we excluded the data from 2 participants in Experiment 1, and no participants in Experiment 2.

Furthermore, in Experiment 1 we excluded trials that used one of the stimulus images from set 1 and one from set 2 because participants performed at chance level with those images in the emotion and in the sex task, respectively. The problem stimulus from set 2 was replaced with another for

Experiment 2. In Experiment 2, due to an experimenter error, we excluded data from one image in the first 13 participants, affecting 2.8% of the total number of trials. Finally, on a trial level, in both experiments, we excluded observations with a response time above or below 3 SDs of each participant’s mean response times across conditions (1.88% of the total number of trials for Experiment 1 and 2.9% for Experiment 2).

Error rate was low across conditions (∼10%) in both experiments (see Supplementary Table 1 and 2). In keeping with previous literature on this task, we focused our analyses on response times in accurate trials. In Experiment 1 we conducted a 2 (Stimulus set: Set 1 or Set 2) x 2 (Task: Sex or Emotion) x 2 (Block: Control or Orthogonal) repeated measures ANOVA on response times in accurate trials. Furthermore, to test which stimulus set in Experiment 1 led to more reliable Garner interference effect, we conducted two follow-up Task x Block ANOVAs, separately for each stimulus set. We also compared the effect sizes of the Garner interference effect (i.e. comparison of orthogonal vs. control block) in the emotion task for the two stimulus sets. In Experiment 2 we conducted a 2 (TMS Stimulation: EBA-V1 or V1-EBA) x 2 (Task: Sex or Emotion) x 2 (Block: Control or Orthogonal) ANOVA, also on response times in accurate trials.

To test for Garner interference (see **Figure 1B**), we followed up the ANOVAs with bi-directional paired t-contrasts directly comparing raw RTs in the orthogonal and control blocks within each task, as well as raw RT for the two tasks in the control condition. Note that using a two-tailed test is a conservative approach, due to the strongly directional hypothesis that performance in Orthogonal blocks should be similar to, or worse than, Control blocks. However, we did not apply a multiple-comparison correction to avoid inflation of type-2 error in the task with least Garner interference. To support interpretation of null results as evidence for parallel processing of sex and emotion, we also report base 10 Bayes factors (BF10). Briefly, BF10 below 1 indicates that the null hypothesis is more likely, and values below 1/3 are generally taken as moderate, reliable evidence (or better) in favour of the null hypothesis (Jeffreys, 1961; Lee & Wagenmakers, 2014; Ly et al., 2016). To further explore possible interactions between sex and emotion idiosyncratic to each stimulus emotion x sex combination, in keeping with recent work of Craig and Lipp (2023), we report in the **Supplementary Materials** reaction time analyses on the simple categorization performance in the orthogonal condition for each task in both experiments.

## 3. Results

### 3.1 Experiment 1

The ANOVA revealed a main effect of Task (F(1,29) = 20.88, *p* < 0.001, η^2^p = 0.42) that was qualified by a significant Task x Block interaction (**Figure 3A**, F(1,29) = 10.33, *p* = 0.003, η^2^p = 0.26). Participants were slower in judging the emotional expression in the orthogonal (M = 688.2, SD = 104.9) vs the control condition (M = 663.8, SD = 109.4, t (29) = 3.56, *p* = 0.001, 95% CI = [9.54, 35.28], d = 0.65, BF10 = 26.4). Conversely, when judging sex, participants’ performance did not reliably differ between the control (M = 634.9, SD = 102.2) and the orthogonal condition (M = 624.1, SD = 94.79, t (29) = -1.17, *p* = 0.25, 95% CI = [-29.79, 8.07], d = -0.21, BF10 = 0.36). This pattern indicates asymmetric Garner Interference: irrelevant variation of sex slowed down emotional expression judgments, but not the converse. No other effect reached significance (all *p*s > 0.13), including the three-level interaction of Stimulus Set x Task x Block (F(1,29) = 1.18, *p* = 0.29, η^2^p = 0.04), suggesting that the overall pattern of asymmetric Garner Interference generalized across the two stimulus sets.

**Figure 3.**
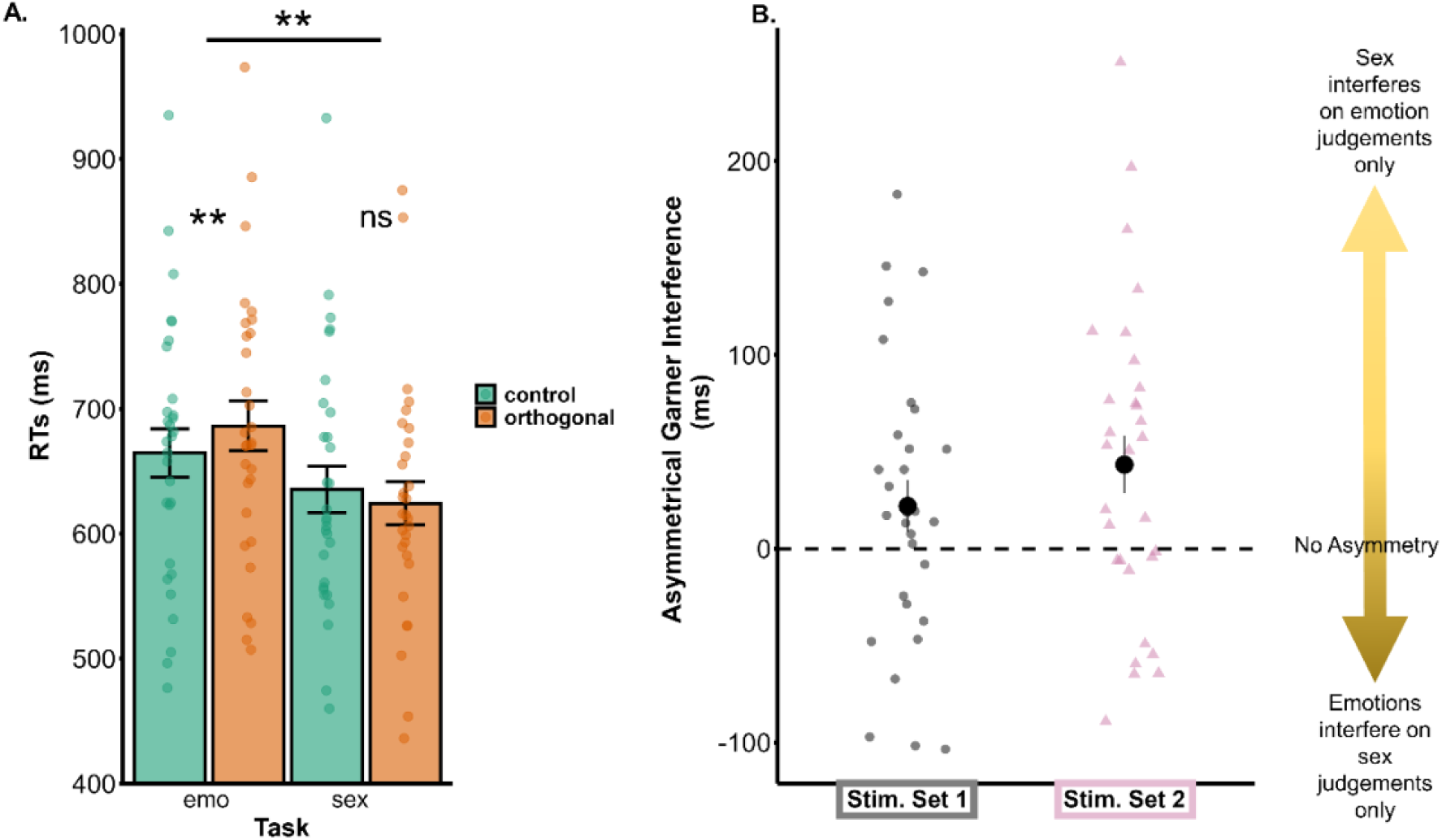
**A.** *Summary of results of Experiment 1, averaging across stimulus sets. The Orthogonal condition led to slower responses in the emotion task but not in the sex task, in which the two blocks did not reliably differ.* **B.** *Pointrange chart showing asymmetrical Garner interference for each stimulus set. This is computed as the difference between two Garner interference effects in the two tasks, each calculated as [mean correct response time in Orthogonal blocks – mean correct response time in Control blocks]. Each point represents data from one participant. Error bars represent the standard error of the mean, * p < 0.05; ** p < 0.01*

For Experiment 2, we wanted to select the stimulus set that would most reliably show asymmetrical Garner interference, to maximize the chance to modulate such interference with TMS. Although no significant three-way interaction between Stimulus set, Task and Block was observed in the main ANOVA, testing the Task x Block interaction separately for each stimulus set showed that the pattern of asymmetrical interference was more consistently observed with stimulus set 2 (See **Figure 3B**, Task x Block interaction: F(1,29) = 8.72, *p* = 0.006, η^2^p = 0.23), as compared to stimulus set 1 where the effect size was smaller (F(1,29) = 2.9, *p* = 0.1, η^2^p = 0.09). With stimulus set 2, Garner interference in the emotion task was also numerically larger (t(29) = 2.76, *p* = 0.010, d = 0.50, mean difference = 25.6 ms, 95 % CI = [6.61, 44.59]) than with stimulus set 1 (t(29) = 2.25, *p* = 0.032, d = 0.41, mean difference = 19.23 ms, 95 % CI = [1.78, 36.68]).

To better illustrate the difference in the Garner interference elicited by the two stimulus sets, we also calculated a Garner interference effect index by subtracting the mean correct RTs in Control block from the mean correct RTs in Orthogonal block for each task and stimulus set. Subtracting the Garner interference index in the sex task from the Garner interference index in the emotion task provided a measure of asymmetrical Garner interference, which was stronger for stimulus set 2 than for stimulus set 1 (Figure 3 B).

Another aspect we considered for selecting the stimulus set to be used in Experiment 2 was to ensure that there were no performance differences between the tasks at baseline (i.e. Control block). One possible account of asymmetric Garner interference is that the two tasks are not matched for general difficulty (Gandolfo and Downing, 2020; Johnstone and Downing, 2017; Garner, 1983; Schweinberger et al., 1999). If the perceptual discriminability of one dimension is stronger than the other, then an asymmetric pattern of interference may be due to these “data limits” rather than to partially integral processing of these body dimensions. To address this possibility, we compared baseline performance for the Control blocks of the two tasks in both stimuli sets. Stimulus set 2 showed no difference between categorization performance at baseline between tasks (t (29) = 0.86, *p* = 0.398, 95% CI = [- 18.81, 45.98], d = 0.16, BF10 = 0.27). Conversely, stimulus set 1 showed differences at baseline in which the sex task (M = 620 ms, SD = 97.8) led to faster response times than the emotion task (M = 664 ms, SD = 114.8, t(29) = 3.69, *p* < 0.001, d = 0.67, 95 % CI = [19.65, 68.44], BF = 35.8).

In sum, we found asymmetrical Garner interference between body sex and fearful/happy body emotions. This asymmetrical interference generalized between two different stimulus sets. However, it was more reliably observed and was not affected by differences in general difficulty between the two tasks at baseline with stimulus set 2, hence that set was adopted for Experiment 2.

### 3.2 Experiment 2

We found a significant TMS stimulation x Task interaction (F(1,28) = 9.61, *p* = 0.004, η^2^p = 0.26), which was further qualified by a significant three-way interaction of TMS stimulation x Task x Block (F(1,28) = 4.69, *p* = 0.039, η^2^p = 0.14) (see **Figure 4**). To follow-up this three-way interaction, we tested the Task x Block interaction for each stimulation session. Following V1-EBA ccPAS, the interaction of Task x Block, indicative of asymmetrical Garner Interference, did not reach significance (F(1,28) = 0.001, *p* = 0.983, η^2^p = 0.00). A Bayes factor t-test showed moderate evidence for the absence of asymmetrical interference (BF10 = 0.20). Indeed, we found no significant Garner interference in the emotion task (Control: M = 563.6, SD = 100.8; Orthogonal: M =575.1, SD = 98.2, t(28) = 1.55, *p* = 0.132, 95 % CI = [-3.7, 26.6], d = 0.29, BF10 = 0.58) nor in the sex task (control: M = 566.8, SD = 99.0; Orthogonal: M =577.9, SD= 85.6, t(28) = 0.98, *p* = 0.337, 95 % CI = [-12.3, 34.6], d = 0.18, BF10 = 0.31).

**Figure 4.**
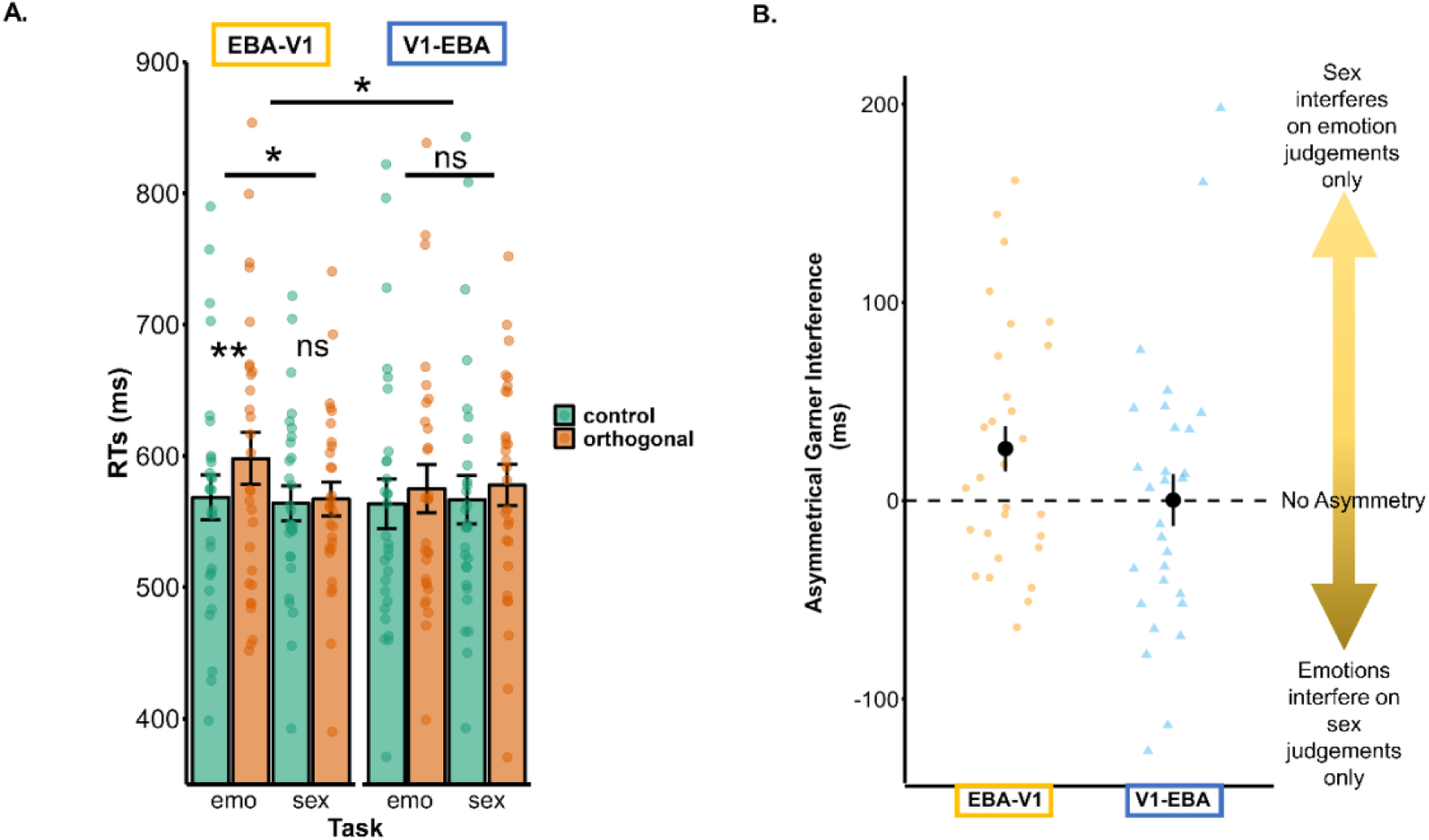
**A.** *Results of Experiment 2. The orthogonal condition led to slower responses in the emotion task but not in the sex task, but this Garner interference effect was significantly modulated by neurostimulation.* **B.** *Pointrange chart showing asymmetrical Garner interference as a function of stimulation direction. Strengthening feedforward connections between V1 and the EBA eliminated the interference that the irrelevant variation of emotional body expressions had on the task-relevant sex judgements. Asymmetrical interference remained intact after strengthening EBA-V1 feedback connections. Error bars represent the standard error of the mean, * p < 0.05; ** p < 0.01*

Conversely, following EBA-V1 stimulation, the interaction of Task x Block was significant (F(1,28) = 5.28, *p* = 0.029, η^2^p = 0.16). Garner interference was present in the emotion task (Control: M = 568.6, SD = 91.5; Orthogonal: M =598, SE = 106.6, t(28) = 2.95, *p* = 0.006, 95 % CI = [9, 49.9], d = 0.55, BF10 = 6.70), but it was not significant in the sex task (Control: M = 564, SD = 71.0; Orthogonal: M = 567.2, SD = 69.3, t(28) = 0.45, *p* = 0.658, 95 % CI = [-11.5, 17.9], d = 0.08, BF10 = 0.22).

To test for a possible difference in performance between the two tasks induced by the stimulation in either block, we tested whether the Control and Orthogonal conditions differed between the two tasks within each stimulation session. In both stimulation conditions, the categorization response times in the Control conditions did not differ significantly (V1-EBA: sex M = 567, SD = 99.0, emotion M = 564, SD = 100.8, t(28) = -0.23 *p* = 0.820, 95% CI = [-31.4, 25.1], d = -0.04, BF10 = 0.20; EBA-V1: sex M = 564, SD = 71.0, emotion M = 569, SD = 91.5, t(28) = 0.34, *p* = 0.734, 95% CI = [-22.8, 32.0], d = 0.06, BF10 = 0.21).

However, differences between tasks were found in the Orthogonal blocks, as found in Experiment 1. After EBA-V1 stimulation, participants showed slower performance in the orthogonal blocks in the emotion task (M = 598.0, SD = 69.3) than in the sex task (M = 567, SD = 69.3, t(28) = 2.45, *p* = 0.021, 95% CI = [5.0, 56.6], d = 0.45, BF10 = 2.46). In contrast, after V1-EBA stimulation this difference did not reach significance (emotion: M = 575, SD = 98.2; sex: M = 577.9, SD = 85.6, t(28) = -0.22, *p* = 0.83, 95% CI = [-30.0, 24.2], d = -0.04, BF10 = 0.20). This would indicate that TMS effects primarily influenced performance in the Orthogonal condition. However, when directly comparing the Orthogonal blocks of the emotion task in the two stimulation conditions (EBA-V1: M = 598, SD = 106.6; V1-EBA: M = 575, SD = 98.2), this difference was not significant (t(28) = -1.94, *p* = 0.063, 95% CI = [-47.2, 1.3], d = -0.4, BF10 = 1.01). This suggests overall that ccPAS TMS influenced the difference between control and orthogonal blocks in this task, rather than solely one or the other block.

Finally, to further inspect whether stimulation of V1-EBA led to a reduction of asymmetric interference, we directly compared the results of Experiments 1 (for stimulus Set 2 only) and Experiment 2.

Specifically, we ran a TMS Stimulation (V1-EBA/NoTMS) x Task (Emotion/Sex) x Block (Control/Orthogonal) mixed-design ANOVA, with TMS Stimulation as a between-participants variable and Task and Block as within-participants variables. We found a significant three-way interaction (F(1,57) = 4.79, *p* = 0.033, η^2^p = 0.08). In contrast, we found no such interaction when we performed a similar analysis comparing the EBA-V1 stimulation condition to the results of Experiment 1 (F(1,57) = 0.87, *p* = 0.36, η^2^p = 0.02).

## 4. General Discussion

Participants could not selectively attend to happy and fearful body emotional expressions and simultaneously fully ignore information about the sex of the depicted individual. Sex is at least partially automatically processed while participants judge emotional expressions. In contrast, sex judgments were relatively unaffected by task-irrelevant variation of emotional expression. This asymmetric interference pattern was not explained by task differences at baseline, because participants were equally fast in judging the sex and emotional expressions in the control conditions (at least for stimulus set 2). This suggests that asymmetric interference genuinely reflects partially integral processing of these two dimensions of the body.

Asymmetric patterns of Garner interference are sometimes explained by a parallel contingent model (Atkinson et al., 2005; Johnstone & Downing, 2017; Schweinberger & Soukup, 1998; Turvey, 1973). The body shape cues that support sex judgements are relatively invariant. Conversely, the postural cues used for emotional expression judgements are relatively changeable aspects of the body, and more variable. As such, it is possible that the most stable visual cues of the body are automatically processed, providing a stable reference for the encoding of emotional expressions, which may differ systematically between the sexes (Kring & Gordon, 1998). This hypothesised interaction between sex- and emotion-related processing may arise “early” at a forward stage of processing, because of involuntary attention capture by visual cues that influence emotion expression judgements. It may also relate to a “late” feedback stage of processing, when the visual features relevant to accomplish the emotional expression task have been selected, and top-down signals related to expectation or to task goals are fed back to visual cortex (Bar, 2004; Summerfield et al., 2006; Summerfield & de Lange, 2014).

To disentangle these possibilities, we sought to modulate the efficiency of feedforward or feedback synaptic connections between the primary visual cortex (V1) and the extrastriate body area (EBA) using ccPAS. If the behavioural interference between sex and emotion affects feedforward computations from early to later stage of processing, then strengthening forward connections with V1-EBA ccPAS should eliminate, or reduce, the Garner interference of sex on emotional expression judgements. Conversely, if the interference affects a feedback stage of processing, EBA-V1 ccPAS should reduce the same interference. We found clear evidence of the former possibility, such that V1-EBA stimulation removed the interfering effects of sex on categorisation of emotion from the body.

Thus, these neuroscientific findings disambiguate the behavioural finding of asymmetric interference, at least as captured in the present task, rooting its cause in feedforward connectivity from relatively low- to higher-level features of the body shape.

A basic assumption of this interpretation is that ccPAS refines the flow of processing along the stimulated pathway, making the perceptual system less vulnerable to the intrusion of task-irrelevant dimensions. An alternative mechanistic interpretation of ccPAS effects could be that boosting feedfoward or feedback connectivity should *increase* the intrusion of the task-irrelevant dimension of sex on emotion processing. This alternative mechanism, however, was not supported by the findings, since we did not observe an increase of Garner interference (compared Experiment 1) after boosting either feedforward or feedback connectivity. We, thus, propose that ccPAS exerted its effects by refining the processing of the task-relevant dimensions.

Our hypothesis is that strengthening synaptic connections from V1 to EBA increased the ability to filter out task irrelevant information, by increasing access to those low-level visual features that are uniquely related to the task at hand. We identify two potential mechanisms that may be behind our neurostimulation findings. First, increasing feed-forward connections may have boosted the bottom-up processing of those low-level visual features that are related to emotional expression judgments and that do not overlap with features that also relate to differences between sexes (cf. Becker et al., 2007). Second, and not mutually exclusively, increasing feed-forward connections may competitively reduce the weight of feedback connections that relate to entangled sex and emotion representations. In other words, if sex provides a top-down reference for emotion judgements (Oldrati et al., 2020), increasing feedforward connectivity might reduce the weight of biased associations of sex with emotions due to top-down influences, leading to reduced interference. This latter possibility, however, is less likely, as it would have also predicted a reduction of the Garner interference following modulation of feedback connections between EBA and V1, relative to the baseline data from Experiment 1. Nevertheless, since we stimulated feedforward and feedback connectivity between V1 and EBA at a fixed time interval based on previous studies of striate-extrastriate connectivity (Romei et al., 2016), we cannot rule out that a different pattern of results may appear at a different timings.

Future studies should explore network-specific timing adjustments to replicate the present effect, and to evaluate the potential for an inverse pattern when targeting feedback connectivity at later interstimulus intervals.

Our analyses focused on accurate reaction times as typically used in Garner selective attention paradigms. This paradigm and its capacity to capture the perceptual dimensionality of complex stimuli is well understood (e.g. Algom and Fitousi, 2016). However, given the high accuracy in the categorization performance (see Supplementary Table 1 and 2), this approach is less suitable for assessing whether the ccPAS stimulation effects found here reflect an improvement in the accuracy of emotion discrimination. Future ccPAS studies could use tasks more focused on accuracy to assess whether strengthening forward V1-EBA connections improves detection of emotional expressions in noise, for example by using signal detection theory measures (Bang & Rahnev, 2017; Tarasi et al., 2022, 2023), or by assessing emotion categorization performance under working memory load (Lavie, 2005; Ramsey et al., 2019).

Asymmetric interference of sex and emotion judgments is not always found in studies using the Garner approach (Gandolfo & Downing, 2020b and Craig & Lipp, 2023). What is the reason for these mixed findings across different studies? To address this, it is useful to look back at the evidence from research on sex and emotion in face perception. For example, Le Gal & Bruce (2002) investigated the relationship between male or female faces expressing either angry or surprised emotions, reporting no evidence for Garner interference in either task, suggesting independence between these two dimensions. Conversely, in another study using fear and happy facial expressions (the emotional expressions we tested here), Atkinson and colleagues (2005) found evidence for asymmetric Garner interference such that sex interfered with the judgments of happy and fearful emotions, but not *vice versa*. More recently, Becker (2017) reported symmetrical interference between sex and angry/happy emotions in face judgments. Yet, the interference of emotion on sex was numerically smaller than *vice versa*, and it disappeared when controlling for general attentional skills of the participants.

These previous face categorisation studies suggest 1) that sex and (at least some) emotional expressions from the face are not fully independently processed, and 2) that interference from invariant (sex) to changeable aspects of the face (emotion) seems to be more robust than the converse. The interference of sex on emotional judgements persists even in absence of baseline differences in the processing speed of the two face characteristics (Atkinson et al., 2005), and after accounting for general cognitive control abilities (Becker, 2017).

In body perception, the same aspects may be considered. Here, we tested for the first time the combination of happy and fearful emotions and their interaction with sex. It is possible that the asymmetric Garner interference reported here is specific to the emotions we have tested. Another possibility, not mutually exclusive, is that interference patterns may in part be idiosyncratic to a given stimulus set. Indeed, the same emotion may be expressed through highly different postures, which may in turn lead to more or less overlap with the features that relate to sex. However, the asymmetric pattern of interference reported here was similar across two different stimulus sets, despite the visual differences e.g. in clothing and intensity of emotional expression. Still, the interference was more reliable in stimulus set 2 (Borgomaneri et al., 2012), which showed bodies with less clothing and, perhaps with more intentional and intense emotional expression than stimulus set 1 (Gandolfo & Downing 2020b). Future research may examine more systematically how interactions between sex and emotion judgements are affected by stimulus details, the specific emotions tested, and other more general dimensions related to emotions, such as valence, intensity, and the relative salience of open versus closed postures (Dael et al., 2012; James, 1890; Kleinsmith & Bianchi-Berthouze, 2007; Poyo Solanas et al., 2020; Rossberg-Gempton & Poole, 1993).

An additional determining factor of Garner interference, besides the variation of the irrelevant dimension as such, relates to higher stimulus uncertainty in the orthogonal compared to the control block (Algom and Fitousi, 2016; Melara and Algom, 2003). It is possible that the interference of sex over emotion judgments observed here was determined by increased stimulus uncertainty and that forward ccPAS reduced the costs of such uncertainty in Experiment 2. However, if the interference was entirely driven by uncertainty, we would have likely observed symmetrical and not asymmetrical interference in both experiments. Indeed, the stimulus uncertainty increased equally in both of the orthogonal blocks tested: the orthogonal block of the emotion task (where we observed interference) and the orthogonal block of the sex task (where the interference was not observed). Human bodies convey several additional meaningful dimensions besides sex and emotion (such as age, race).

Future Garner studies can include additional irrelevant variation of another body dimension (a “correlated filtering” condition – Burns, 2016), for example the age, to disentangle more directly the behavioural cost caused by stimulus uncertainty on top of the one caused by irrelevant stimulus dimensions as such.

## 5. Conclusions

Our findings add to our understanding of the structure of visual representation of the human body, suggesting that body sex information is processed integrally with the postural information conveying happy and fearful emotions. We provide novel neuroscientific evidence that the interactions between those dimensions revealed by Garner tasks likely emerge as a result of forward processing, during the perceptual analysis of the stimuli, rather than as a result of top-down links between the social categories. Finally, our findings confirm that ccPAS is a promising TMS protocol for investigating of bottom-up and top-down processes in visual cognition and person perception.

^1^ In line with current guidance (i.e. Sager Guidelines - https://researchintegrityjournal.biomedcentral.com/articles/10.1186/s41073-016-0007-6/tables/1) we use “sex” to refer to a categorical distinction between body images and “gender” in reference to our participants. We regard “sex” as the appropriate term for our experimental manipulation of body shape, as these images are presented without any further context that would be likely to generate a gender interpretation.

## Author Contribution

**Marco Gandolfo:** Conceptualization, Data Curation, Formal Analysis, Data Visualization, Writing – Original Draft, Writing – Review & Editing, Investigation, Funding Acquisition.

**Giulia D’Argenio:** Conceptualization, Investigation.

**Paul E. Downing:** Conceptualization, Writing – Original draft, Writing – review & editing.

**Cosimo Urgesi:** Conceptualization, Funding acquisition, Writing – review & editing, Supervision, Funding Acquisition.

## Data and code availability statement

The raw and processed data generated from this project are available at the Open Science Framework repository alongside the analysis code - https://osf.io/mtxfw/. The Stimuli from Gandolfo and Downing (2020), including those used in Experiment 1 were also made publicly available in the same repository.

## Acknowledgements

We thank Marco Zanon for providing scripts for controlling the TMS machine; and Sonia Betti and Alessandra Finisguerra for help with the TMS setup.

