## Supplementary Material for "Boosting forward connectivity between primary visual and body selective cortex reduces interference between sex and emotion judgements of bodies"

**Supplementary Materials**

| Version | task | realblock | Accuracy | SD |
|---|---|---|---|---|
| **Stim. Set 1** | Emotion | control | 0.89 | 0.05 |
| **Stim. Set 1** | Emotion | orthogonal | 0.9 | 0.07 |
| **Stim. Set 1** | Sex | control | 0.9 | 0.06 |
| **Stim. Set 1** | Sex | orthogonal | 0.9 | 0.07 |
| **Stim. Set 2** | Emotion | control | 0.92 | 0.04 |
| **Stim. Set 2** | Emotion | orthogonal | 0.91 | 0.05 |
| **Stim. Set 2** | Sex | control | 0.89 | 0.07 |
| **Stim. Set 2** | Sex | orthogonal | 0.9 | 0.06 |

**Supplementary Table 1.** *Mean accuracy and standard deviation in each condition across participants in Experiment 1.*

| Version | Task | Block | Accuracy | SD |
|---|---|---|---|---|
| **EBA-V1** | emotion | control | 0.91 | 0.05 |
| **EBA-V1** | emotion | orthogonal | 0.92 | 0.05 |
| **EBA-V1** | sex | control | 0.9 | 0.05 |
| **EBA-V1** | sex | orthogonal | 0.91 | 0.05 |
| **V1-EBA** | emotion | control | 0.92 | 0.04 |
| **V1-EBA** | emotion | orthogonal | 0.92 | 0.05 |
| **V1-EBA** | sex | control | 0.89 | 0.08 |
| **V1-EBA** | sex | orthogonal | 0.9 | 0.07 |

**Supplementary Table 2**. *Mean accuracy and standard deviation in each condition across participants in Experiment 2.*

### Analyses on simple categorization performance in the orthogonal blocks

This supplemental analysis follows the logic of previous studies focused on interactions between sex and emotions in body perception (Bijlstra et al., 2019; and Craig and Lipp, 2023). Here, response times and accuracy were analysed in the orthogonal blocks to determine how sex and emotion interact with each other. Differences in categorization performance depending on specific sex-emotion combinations have often been taken as further evidence of interactive processing (or “crosstalk” between dimensions; Plant et al., 2000; Hugenberg and Sczesny, 2006; Bijlstra et al., 2019; Huang et al., 2020; Craig and Lipp, 2023). Particularly, Bijlstra et al., 2019 and Huang et al., 2020 report a categorization advantage for happy emotions in female vs male bodies in an emotion categorization task including emotions with different valence (Angry vs Happy). For sex categorization instead, Craig and Lipp (2023) report faster judgements of angry bodies expressions of males than females. Accordingly, in our study we could expect participants to have better performance in the emotion task for categorizing happiness in female than male bodies, and for categorizing fearful in male than female bodies. In the sex task, participants may show increased performance for fearful male bodies over female bodies.

#### Experiment 1

We ran a Stimulus Set (Set 1/Set 2) x Task (Emotion/Sex) x Stimulus Emotion (Happy/Fearful) x Stimulus Sex (Male/Female) ANOVA on both accurate RTs and mean categorization accuracy of the orthogonal block. On both measures, we observed a significant 4-way interaction (RTs: $F(1,29) = 15.38$, $p < 0.001$, $p\eta^2 = 0.35$; Accuracy: $F(1,29) = 15.02$, $p < 0.001$, $p\eta^2 = 0.34$).

We followed this interaction by splitting by considering each stimulus set separately. Where an interaction between Task x Stimulus Emotion x Stimulus Sex is present, we further broke this result down by analysing the categorization performance in each task separately, following the approach of Craig and Lipp, 2023.

*Stimulus Set 1*

With stimulus set 1 we observed no interaction of Task x Stimulus Sex x Stimulus emotion (RT - $F(1,29) = 0.001$, $p = 0.975$, $p\eta^2 < 0.01$; Accuracy - $F(1,29) = 0.47$, $p = 0.499$, $p\eta^2 = 0.16$).

In the RT analysis we observed a main effect of task $F(1,29) = 28.18$, $p < 0.001$, $p\eta^2 = 0.98$), with the sex task (M = 617ms, SE = 16.19) being overall faster than the emotion task (684ms, SE = 21.42), a main effect of emotion $F(1,29) = 5.77$, $p = 0.023$, $p\eta^2 = 0.17$), with fearful stimuli being categorized faster (640ms, SE = 16.63) than happy stimuli (661ms, SE = 20.09). The two main effects were qualified by a Task x Emotion interaction ($F(1,29) = 6.97$, $p = 0.013$, $p\eta^2 = 0.19$). In the emotion task, participants were faster in categorizing fearful than happy stimuli ($t(29) = 2.69$, $p = 0.012$, $d = 0.52$). This difference was not significant in the sex task, when emotional expressions were irrelevant ($t(29) = 0.28$, $p = 0.96$, $d < 0.01$). No other effect reached significance (all *p*s > 0.13).

In the accuracy analysis we only observed a Task x Stimulus Sex interaction ($F(1,29) = 10.22$, $p = 0.003$, $p\eta^2 = 0.26$). In the sex task, female stimuli (M = 0.87, SE = 0.02) were judged less accurately than male stimuli (M = 0.93, SE = 0.02, $t(29) = 2.67$, $p = 0.012$, $d = 0.47$). This was not the case when the sex was task irrelevant, during the emotion categorization task ($t(29) = -1.36$, $p = 0.19$, $d = -0.13$). No other effect reached significance (all *p*s > 0.08).

Together, using a simple categorization approach we find little evidence for "crosstalk" between sex and emotion categorization with stimulus set 1.

*Stimulus Set 2*

With Stimulus Set 2, we observed a Task x Stimulus Emotion x Stimulus Sex interaction on both accurate RTs ($F(1,29) = 37.78$, $p < 0.001$, $p\eta^2 = 0.57$) and accuracy scores ($F(1,29) = 23.33$, $p < 0.001$, $p\eta^2 = 0.45$). To follow-up on this 3-way interaction we then split the design by task and assessed whether a Stimulus Emotion x Stimulus Sex interaction was observed. Results of the accurate RTs are shown in **Figure S1** below.

In the emotion task, we observed a Stimulus Emotion x Stimulus Sex interaction on both accurate RTs ($F(1,29) = 14.46$, $p < 0.001$, $p\eta^2 = 0.33$) and accuracy scores ($F(1,29) =$

23.13, $p < 0.001$, $p\eta^2 = 0.44$). Participants were faster in judging happy (M = 662ms, SE = 23) than fearful females (M = 694ms, SE = 23.05), t(29) = -2.21, $p = 0.035$, d = -0.44) and faster in judging fearful (M = 685ms, SE = 21.39) than happy males (M = 722ms, SE = 25.23, t(29) = 2.64, $p = 0.013$, d = 0.49). In addition, responses to happy bodies were faster when the stimulus was a female (M = 661ms, SE = 18.70) than a male (M = 722ms, SE = 25.23, t(29) = 4.55, $p < 0.001$, d = 0.83).

The same pattern was observed in accuracy, with participants being better at categorizing happy (M = 0.94, SE = 0.01) vs fearful females (M = 0.90, SE = 0.01, t(29) = 2.56, $p = 0.016$, d = 0.40) and better at categorizing fearful (M = 0.94, SE = 0.01) vs happy males (M = 0.86, SE = 0.02, t(29) = -3.81, $p < 0.001$, d = -0.83). Responses to happy bodies were more accurate when the stimulus was a female (M = 0.94, SE = 0.01) than a male (M = 0.86, SE = 0.02, t(29) = -4.03, $p < 0.001$, d = 0.86). Conversely, responses to fearful bodies were more accurate for male (M = 0.93, SE = 0.01) vs female bodies (M = 0.90, SE = 0.01, t(29) = 2.97, $p = 0.006$, d = 0.37).

In the sex task, we also observed an Emotion x Stimulus Sex interaction on both RTs ($F(1,29) = 14.09$, $p < 0.001$, $p\eta^2 = 0.33$) and accuracy scores ($F(1,29) = 5.39$, $p = 0.027$, $p\eta^2 = 0.16$). Participants were faster in judging males (M = 599ms, SE = 17.87) than females (M = 647ms, SE = 24.22) when bodies were expressing happy emotions, t(29) = -3.28, $p = 0.003$, d = -0.70) while there was no significant difference in sex categorization for fearful males (M = 651ms, SE = 22.57) vs fearful females (M = 639ms, SE = 22.69, t(29) = 0.97, $p = 0.341$, d = 0.18). In addition, responses to male bodies were faster when the stimulus was happy (M = 599ms, SE = 17.87) than fearful (M = 651ms, SE = 22.56, t(29) = -4.43, $p < 0.001$, d = 0.77) and there was no reliable difference between happy vs fearful in categorization of female bodies (t(29) = -.85, $p = 0.40$, d = 0.11).

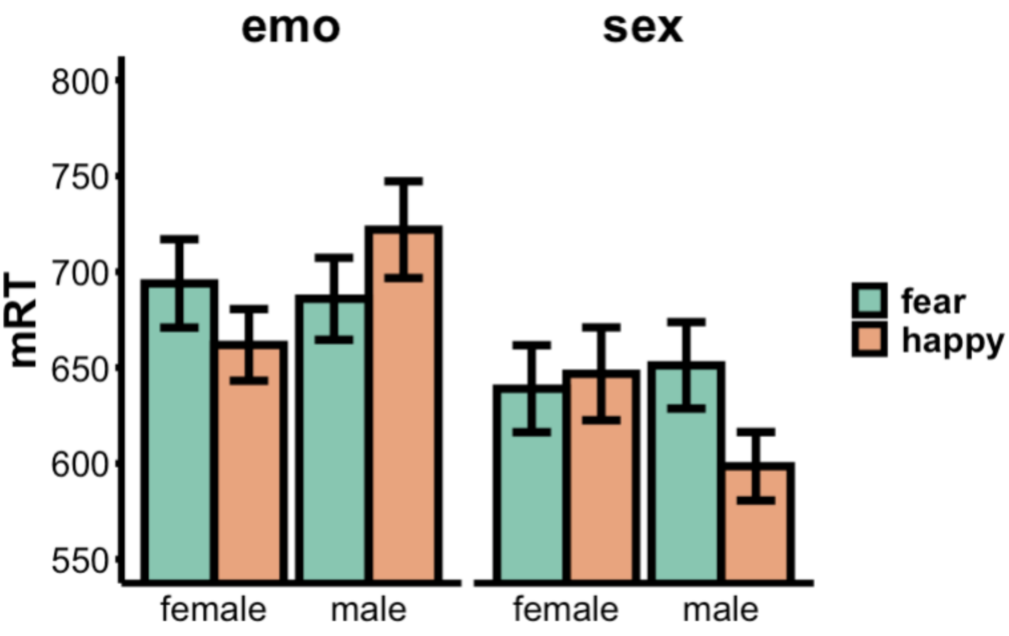

*Figure S1. Simple categorization performance in reaction times in accurate trials (y axis) as a function of stimulus emotion (X axis) and stimulus sex (Coloured legend) in each task. Here we only present data from Stimulus Set 2, in which differences in simple categorization performance were found.*

Together, the simple categorization analysis of Experiment 1 shows that in the emotion task, particularly with stimulus set 2 (which also showed the Garner Interference more reliably), categorization performance of emotion is influenced by stimulus sex in line with previous reports. That is, happy bodies were categorized better and faster when the stimulus observed was a female than a male. Conversely, fearful bodies were categorized better, and faster, when males. The same relationship, however, didn't hold in the sex task, in which, contrary to the expectation coming from social categorization studies (e.g. Craig and Lipp, 2023), male bodies were detected better and faster when showing an happy emotion. Most of the literature on simple categorization tasks studying the relationship of sex and emotion used emotion categorization tasks (Hugenberg and Sczezny, 2006; Craig and Lipp, 2018; Bijlstra et al., 2019) rather than sex tasks. It is possible that in a sex tasks absolute differences in categorization performance due to emotional body postures are better explained by which features a given emotional body posture makes visible, and therefore useful for categorization (cf. Martin and Macrae, 2007; Gandolfo and Downing, 2020). Alternatively, a violation of expectation (i.e. an effect of surprise of seeing an happy male) may explain faster reaction times in the answers to males when showing an happy emotion.

**Experiment 2**

In Experiment 2, we repeated the same simple categorization approach analysis to assess whether ccPAS stimulation affected categorization of certain combinations of sex and emotion, in addition to reducing the asymmetric Garner interference as demonstrated in the main manuscript.

We therefore tested categorization performance in reaction times and accuracy in a Stimulation (EBA-V1/V1-EBA) x Task (Emotion/Sex) x Stimulus Emotion (Happy/Fear) x Stimulus Sex (Male/Female). In both analyses, we found no significant 4-way interaction (RT: $F(1,28) = 0.172$, $p = 0.681$, $p\eta^2 < 0.01$; Accuracy: $F(1,28) < 0.01$, $p = 0.956$, $p\eta^2 < 0.01$). Instead, in the reaction times we observed a Task x Stimulus Emotion x Stimulus Sex interaction ($F(1,28) = 25.14$, $p < 0.001$, $p\eta^2 = 0.47$) which did not reach significance in the accuracy performance ($F(1,28) = 3.25$, $p = 0.084$, $p\eta^2 = 0.10$). The reaction time results are shown in **Figure S2**.

For reaction times, the Stimulus Emotion x Stimulus Sex interaction was significant in the sex task ($F(1,28) = 14.68$, $p < 0.001$, $p\eta^2 = 0.34$) and marginally significant in the emotion task ($F(1,28) = 3.67$, $p < 0.066$, $p\eta^2 = 0.12$). In both tasks (see Figure S2) the pattern of categorization responses was very similar to Experiment 1, with emotion categorization being faster in happy male vs happy females ($t(28) = 2.41$, $p = 0.023$, d = 0.45) and sex categorization being faster for happy than fearful male ($t(28) = =5.49$, $p < 0.001$, d = 0.77).

Together, it is clear that ccPAS stimulation did not affect categorization performance as such, here in the sense of modulating existing "crosstalk" links between specific levels (e.g. female and happy). Rather, stimulation appears to influence the attentional interference of sex on emotion that accumulates over the orthogonal vs the control blocks. This suggests that the TMS effects of ccPAS affected the perceptual processing of the two dimensions (the whole stimulus) rather than categorization of the individual dimensions per se.

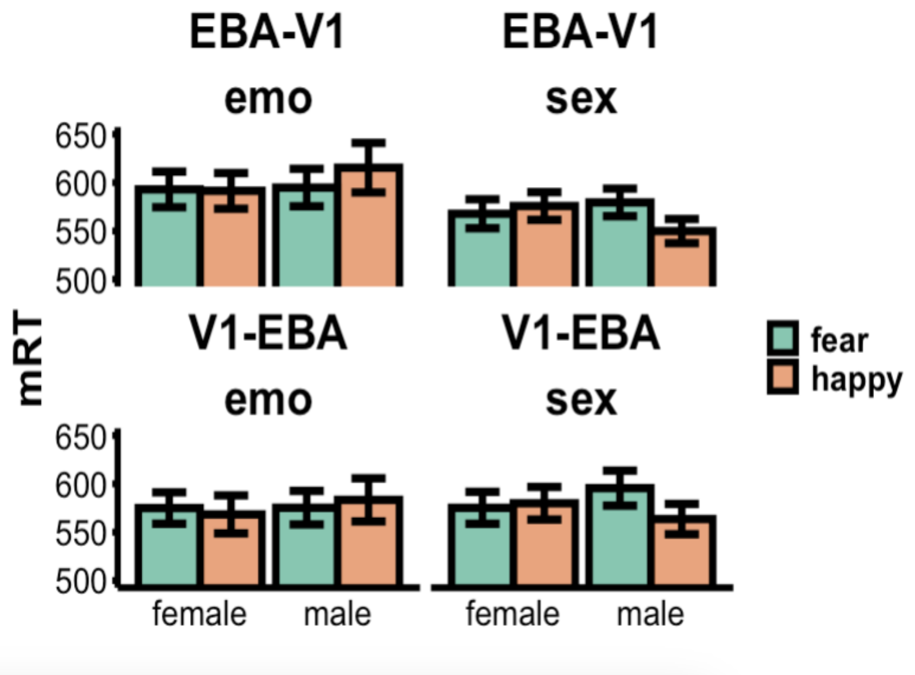


*Figure S2. Reaction times of simple categorization performance as a function of Stimulation Session, Task, stimulus Sex and Stimulus Emotion. No effect of stimulation was observed on simple categorization performance.*
